# Environmental Triggers of *lrgA* Expression in *Streptococcus mutans*

**DOI:** 10.1101/668731

**Authors:** Ivan Ishkov, Sang-Joon Ahn, Kelly C. Rice, Stephen J. Hagen

## Abstract

The *lrgAB* and *cidAB* operons of *Streptococcus mutans* encode proteins that are structurally similar to the bacteriophage lambda family of holin-antiholin proteins, which are believed to facilitate cell death in other bacterial species. Although their precise function is not known, *cidAB* and *lrgAB* are linked to multiple virulence traits of *S. mutans*, including oxidative stress tolerance, biofilm formation, and autolysis. The regulation of *cidAB* and *lrgAB* is still not understood, as these operons show opposite patterns of expression as well as a complex dependence on growth conditions. We have used a microfluidic approach, together with single-cell imaging of a fluorescent gene reporter, to identify with greater precision the cues that trigger *lrgA* expression and characterize cell-to-cell heterogeneity in *lrgA* activity. *lrgA* activates very abruptly at stationary phase, with a high degree of synchrony across the population. We find this activation is controlled by a small number of inputs that are sensitive to growth phase: Extracellular pyruvate, glucose, and molecular oxygen. Further, activation of *lrgA* appears to be self-limiting, so that *lrgA* is strongly expressed only for a very short interval of time. Consequently, *lrgA* is programmed to switch on briefly at the end of exponential growth, as glucose and molecular oxygen are exhausted and extracellular pyruvate is available. Our findings are consistent with studies showing that homologs of *lrgAB* are linked, together with *lytST*, to the reimport of pyruvate for anaerobic fermentative growth.

**Importance:** The function and regulation of *cidAB* and *lrgAB* in Streptococcus mutans is not understood, although these operons have been clearly linked to stress responses and they show a complex dependence on environmental inputs and growth phase. Identifying specific environmental cues that trigger activation of *lrgAB* has been difficult owing to the cells’ own modification of key inputs such as glucose and oxygen: In *S. mutans* the *lrgAB* operon is strongly upregulated at the end of exponential phase, where growth conditions in a bulk culture become poorly defined. Here we have used microfluidics to apply precise control of environmental inputs to *S. mutans* and identify specific chemical cues that activate *lrgAB*. We find that rigorously anaerobic conditions and the presence of extracellular pyruvate are sufficient to induce *lrgAB* expression, suggesting that *lrgAB* is timed to activate just as pyruvate fermentation becomes favorable.

## Introduction

The oral pathogen *Streptococcus mutans* (1) possesses two operons designated *cidAB* (SMU.1701/1700) and *lrgAB* (SMU.575/574) (2), which are closely homologous to the *lrgAB* and *cidAB* operons that have been extensively studied in organisms such as *Bacillus subtilis* and *Staphylococcus aureus* (3–10). Sequence homology indicates that *cidAB* and *lrgAB* encode membrane proteins that are similar to holin-antiholin membrane proteins of the bacteriophage lambda family (11–14), which control autolysis and cell death by modulating the permeability of the bacterial cell wall (10, 12, 13, 15, 16). In *S. mutans*, deletions in *cidAB* or *lrgAB* have been shown to affect virulence-related behaviors such as autolysis, genetic competence, antibiotic resistance, biofilm development, and response to heat and oxidative stresses (11, 17–20). Consequently *lrgAB* and *cidAB* have been viewed as potentially encoding an *S. mutans* holin-antiholin system that responds to conditions of environmental stress by triggering autolysis and cell death (11, 21). However the regulation of *cidAB* and *lrgAB* in *S. mutans* is complex, and the precise function of these genes has not yet been established (11, 17, 18). Expression of *S. mutans cidAB* and *lrgAB* appears linked to several two component signal transduction systems and to carbon catabolite repression, and these two operons display opposite patterns of expression during growth and maturation of a culture (11, 17, 18, 22). The link to variable parameters such as carbohydrate and growth phase has made it difficult to identify specific cues that control the timing and extent of *lrgAB* and *cidAB* transcription. In addition, the kinetics and population heterogeneity of *S. mutans lrgAB* and *cidAB* expression have not been investigated. In this work we apply microfluidic and single-cell approaches to better define the environmental inputs and identify cues that control *lrgAB*. We also characterize the temporal profile and cell-to-cell heterogeneity of the *lrgAB* response to these cues.

In *S. mutans* the *cid* operon consists of *cidA* (342 bp) and *cidB* (696 bp), which overlap by 4 nucleotides (11). The *lrg* operon includes *lrgA* (468 bp) and *lrgB* (732 bp) (11). Both *cidAB* and *lrgAB* are sensitive to glucose availability, although the two operons behave oppositely. When *S. mutans* grows in limited glucose (less than 20 mM), *lrgAB* is not strongly expressed until the onset of stationary phase (11, 22). Higher initial glucose concentrations, exceeding 20 mM, reduce the stationary phase expression of *lrgAB*. By contrast, *cidAB* is robustly expressed during early growth in high glucose concentrations, but is much less active later in growth or when initial glucose concentrations are less than about 20 mM (11, 22). Kim et. al. have recently identified a catabolite responsive element (*cre*-site) region in the promoters of *cidAB* and *lrgAB*, indicating that the catabolite repression protein CcpA may enhance or suppress *cidAB* and *lrgAB* expression during early and late growth stages respectively (22).

Several studies have found that *cidAB* and *lrgAB* respond to molecular oxygen, and that deletions in either operon affect the ability of *S. mutans* to tolerate oxidative stress (2, 11). The Δ*lrgAB* and Δ*cidAB* deletion strains did not grow under aerobic conditions, although their anaerobic growth was reported similar to wild type (11). Similarly, Δ*lrgAB* and Δ*cidAB* strains were unusually sensitive to superoxide anion (generated by paraquat) although not to hydroxyl radical (generated by hydrogen peroxide) (11). Microarray experiments indicated that *lrgA* transcription increased in the presence of molecular oxygen during exponential growth phase (2). A transcriptional profiling study found that *lrgAB* transcription at 0.4 OD in a culture grown aerobically was 11-fold higher than in a culture grown in an anaerobic chamber (2). Furthermore, the deletion of *vicK* (18), which through its PAS domain may play the role of a redox sensor in the VicRK two component system, was found to suppress the late-growth onset of *lrgAB* expression (18, 23–25). *lrgA* and *lrgB* were also upregulated in thicker biofilms, perhaps suggesting sensitivity to oxygen conditions or other environmental stresses within the biofilm (6, 26–28).

The LytST two component system also plays a role in *lrgAB* regulation in *S. mutans*, in which the *lytST* operon is located 175 nucleotides upstream of *lrgAB* (11). LytST and its homologs have been closely linked to regulation of *lrgAB* homologs in many bacteria, including *Bacillus* and *Staphyloccus* species as well as *S. mutans* (3, 6, 8, 11, 17, 29). In *S. mutans* the deletion of *lytST* or *lytS* reduced *lrgAB* expression throughout the growth curve and either eliminated (11) or sharply suppressed (17) the 10^3^-10^4^ fold increase in *lrgAB* mRNA levels that occurs late in the growth curve under low glucose conditions (11). This modulation of *lrgAB* induction by *lytS* was slightly greater at low oxygen conditions (17), possibly indicating a link between LytST and environmental oxygen in regulation of *lrgAB*.

These prior findings show that growth-phase sensitive parameters such as glucose and oxygen interact to regulate *lrgAB* and may contribute to the suppression of *lrgAB* until the onset of stationary phase. Understanding this regulation in detail requires a greater degree of environmental control than is achieved through conventional, bulk culture methods. For this reason, we have used a microfluidic approach to maintain precise control of the environmental inputs that are suspected to influence *S. mutans lrgAB*, and to explore the population profile and kinetics of *lrgAB* expression at the individual cell level. By imaging and quantifying activity of a green fluorescent protein reporter for the *lrgAB* promoter in individual *S. mutans* under controlled flow conditions, we are able to identify the environmental inputs that trigger activation of *lrgAB*.

## Results

### A burst of lrgA activity coincides with the onset of stationary phase

To test our P*lrgA-gfp* fluorescent reporter strain and characterize *lrgA* expression in static cultures, we monitored the optical density and fluorescence of the reporter strain growing in well plates containing defined medium (FMC (30, 31)) that was prepared with different initial concentrations of glucose. Figures 1A and 1B show growth curves for the UA159 background and the P*lrgA*-*gfp* reporting strain respectively, growing anaerobically under a layer of mineral oil. Figures 1C and 1D show the green fluorescence (485 nm excitation, 528 nm emission) of the two strains. For both strains, the growth medium contributes a large background fluorescence that declines steadily as the culture grows. In *PlrgA-gfp* however, the green fluorescence increases abruptly above background as the culture enters stationary phase (arrows in Figure 1D), signaling a strong burst of *lrgA* expression. This rapid rise in green fluorescence is transient, as the green fluorescence at later times declines slowly, like that of the UA159 background. The brief duration of the *lrgA* expression burst is apparent from the time-derivative of the fluorescence signal. Comparing Figures 1E and 1F shows that the fluorescent reporter for *lrgAB* is activated for no more than 30-50 minutes at the onset of stationary phase.

**Figure 1:**
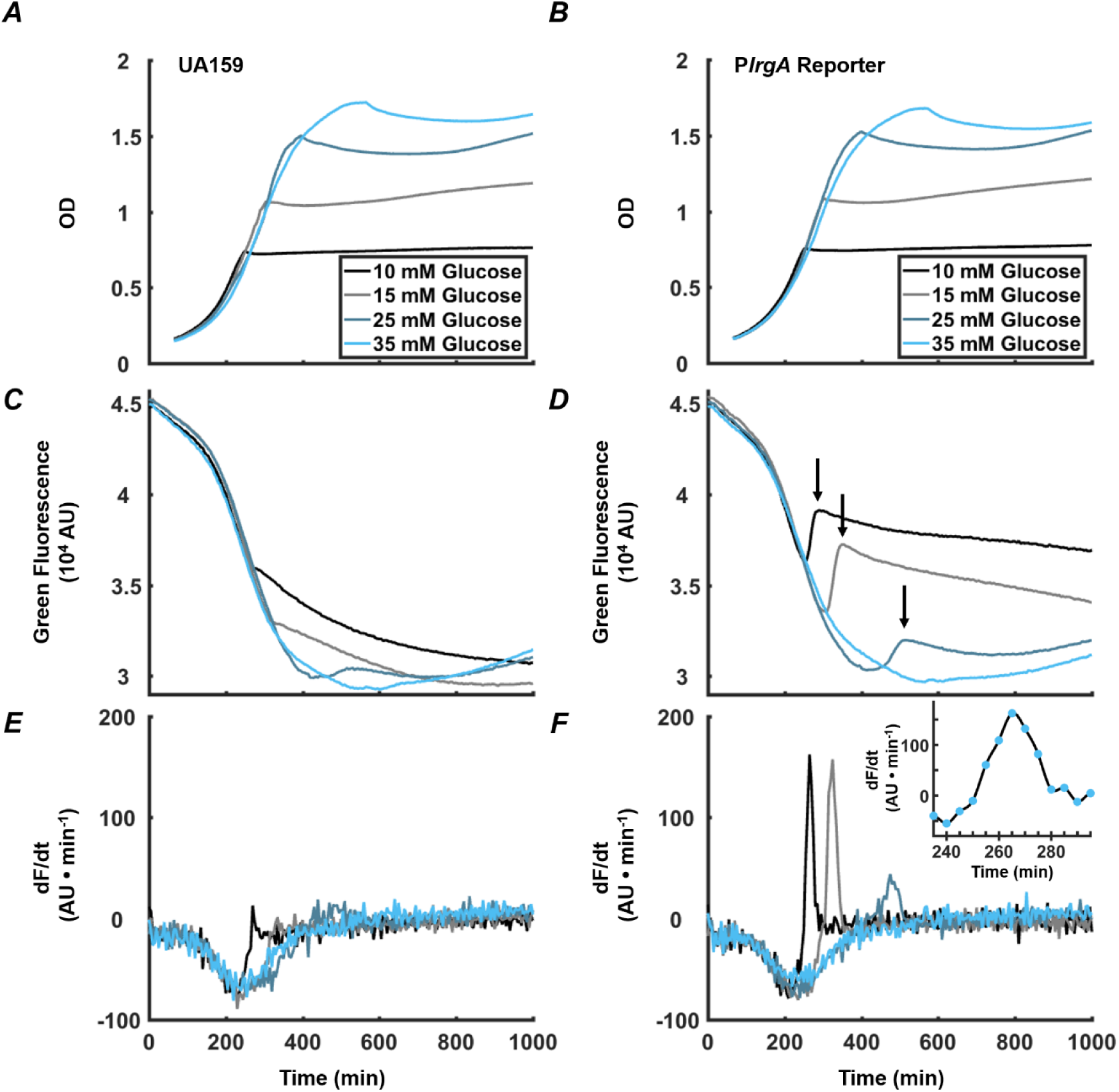
Observation of P*lrgA*-*gfp* fluorescence at the onset of stationary phase in *S. mutans*. Optical density of (**A**) UA159 background and (**B**) P*lrgA-gfp* strain growing in defined medium. Green fluorescence of (**C**) UA159 and (**D)** P*lrgA-gfp* cultures is dominated by the steadily declining fluorescence of the medium, until about 250-300 minutes. The black arrows in (**D**) mark the abrupt burst of fluorescence in the P*lrgA-gfp* strain at the onset of stationary phase. Comparison of the time derivatives of the green fluorescence for (**E**) UA159 and (**F**) P*lrgA-gfp* shows that the burst of *lrgA* expression has a duration of 30-50 minutes. The inset in (**F**) shows the time derivative of reporter fluorescence inh 10 mM glucose.

Figure 1D also shows that the initial glucose concentration of the medium influences the overall amount of *lrgA* expression that occurs during the burst. The size of the fluorescence rise in Figure 1D increases as the initial glucose is raised from 10 mM to 15 mM, but declines as the initial glucose is further raised to 25 mM. At 35 mM initial glucose, the burst is not detected. These data are consistent with transcriptional data showing that *lrgAB* is upregulated 10^3^-10^4^ fold in late exponential phase, relative to early or mid-exponential phase (11), and that very high initial glucose concentrations suppress this upregulation (11, 22).

### The burst of lrgA expression is observed only under anaerobic conditions

Prior studies have found interplay between *lrgAB* expression and molecular oxygen or oxidative stresses (11, 17, 18, 23–25). To more carefully assess the relationship between aerobic or anaerobic conditions and glucose availability on *lrgAB*, we measured the size of the stationary phase burst of reporter fluorescence in well plates that were growing anaerobically (with a mineral oil layer) or aerobically (open to air), with different glucose concentrations. Figure 2 shows that, under anaerobic conditions, increasing the initial glucose to about 10 mM increases the amplitude of the *lrgA* expression burst. However, this amplitude falls monotonically if initial glucose is further increased. In P*lrgA-gfp* cultures grown aerobically, we observed no burst of *lrgA* expression at any initial glucose concentration. Therefore, the burst of *lrgA* expression that occurs in a static culture requires anaerobic conditions as well as a moderately low initial glucose concentration. However, lower glucose concentration does not ensure higher *lrgA* expression; Figure 2C shows that the amplitude of the fluorescence burst declines at initial glucose concentrations smaller than about 10 mM.

**Figure 2:**
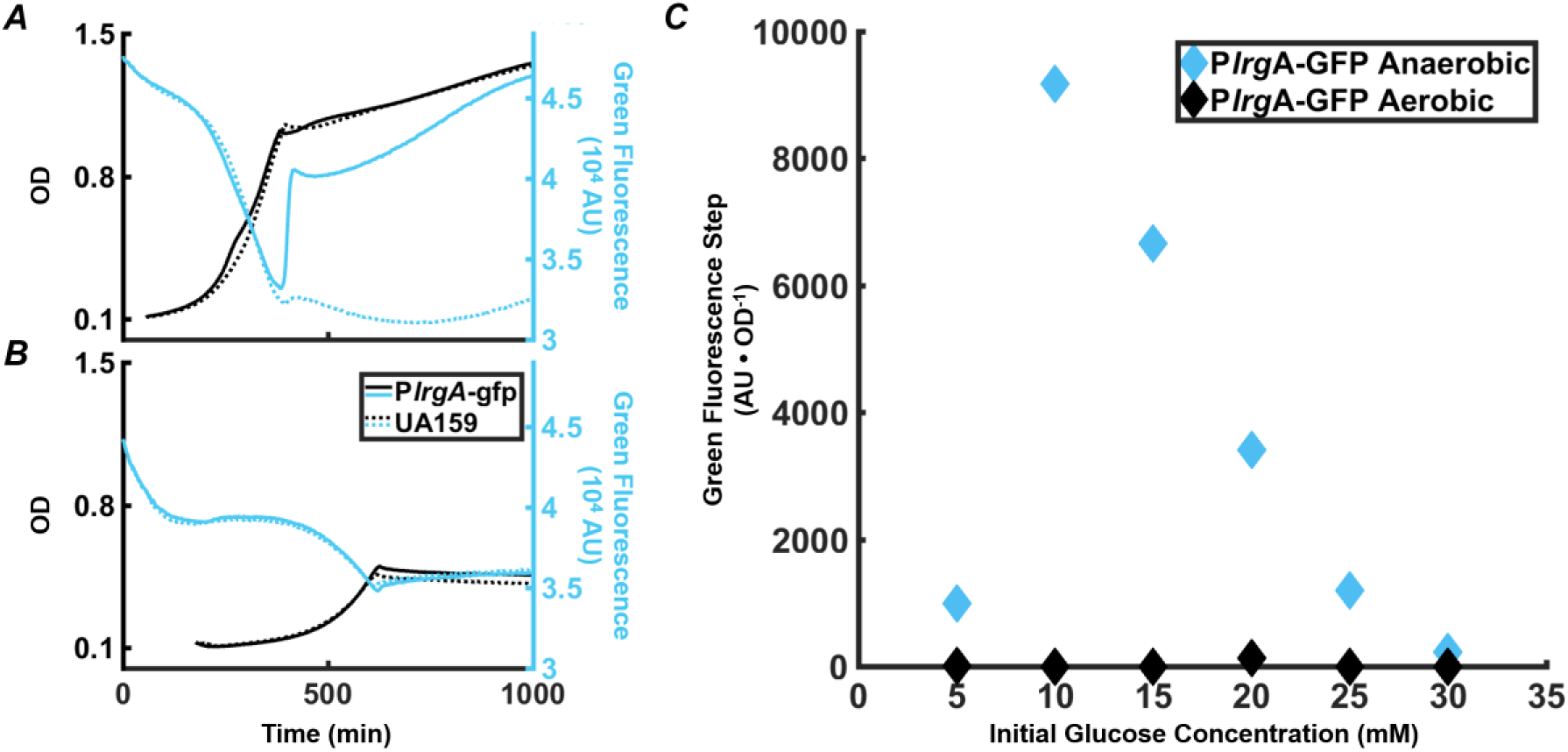
Effect of anaerobic (**A**) and aerobic (**B**) growth on the step increase in *lrgA* activity at stationary phase in static cultures of the *PlrgA-gfp* strain. OD (black curves) and green fluorescence (blue curves) are shown for the reporter (solid curve) and UA159 (dashed curve) strains growing in medium supplemented with 15 mM glucose. (**C**) Comparison of the step in *lrgA* activity for aerobic and anaerobic growth, versus initial glucose concentration. The increase in *lrgA* activity in the reporter strain is measured as the magnitude of the fluorescence step (black arrows in Figure 1D) above background, normalized to the optical density. No fluorescence step is detected in the aerobic cultures.

### Extracellular pyruvate affects stationary phase expression of lrgA

Recent findings that the LytST family of two component systems, which modulate the expression of *lrgA* homologs, can bind and sense external pyruvate (8, 10, 32), and the observation that the pyruvate dehydrogenase complex in *S. mutans* is upregulated in late exponential phase (22), suggest that late growth expression of *lrgAB* in *S. mutans* may be connected to the presence of external pyruvate. We monitored the P*lrgA-gfp* reporter strain growing anaerobically in defined medium to which different concentrations of initial glucose and pyruvate were added. Figure 3 shows that very low concentrations of pyruvate (0 – 0.1 mM) had little effect on the magnitude of the step increase in GFP fluorescence at the onset of stationary phase, regardless of glucose concentration. However, further increases in pyruvate to 1.5 – 8 mM generally enhanced the stationary phase response of *lrgAB*, especially for cells growing at low glucose, 15 mM or less. Higher levels of pyruvate sharply reduced the activation of *lrgA*, until the fluorescence burst became undetectable at 100 mM pyruvate. These data show that initial glucose and pyruvate concentrations constitute a pair of external inputs that can modulate and maximize the stationary phase burst in *lrgA*, although both are inhibitory at higher concentrations.

**Figure 3:**
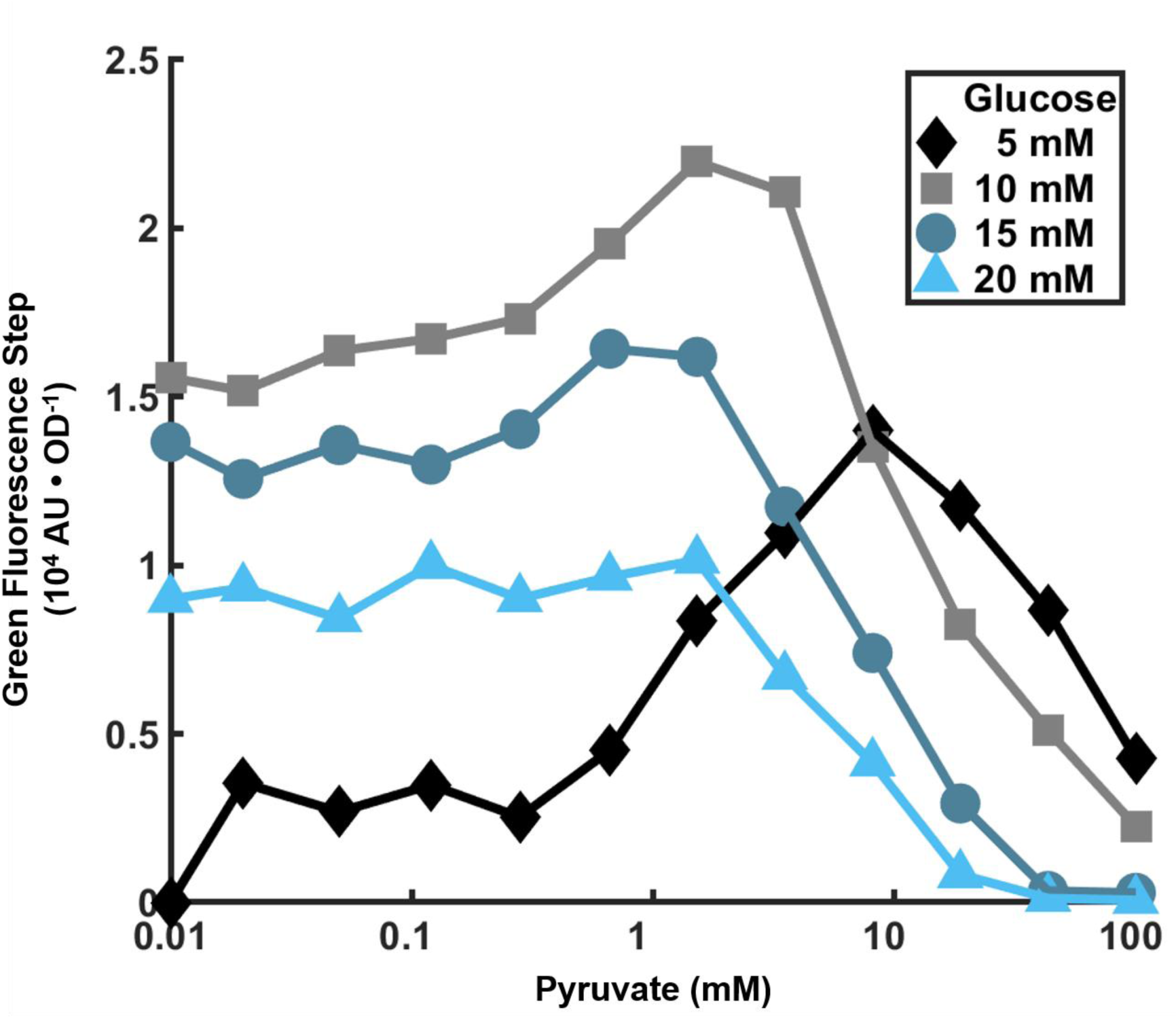
Dependence of *lrgA* expression step on initial pyruvate and glucose concentrations in cultures grown anaerobically. The increment in *PlrgA*-*gfp* reporter fluorescence (relative to baseline) at the onset of stationary phase is shown, normalized to the optical density.

### Expression of lrgA in bulk cultures at stationary phase is heterogeneous

The very rapid burst of P*lrgA-gfp* fluorescence in Figure 1E shows that the timing of *lrgAB* activation is highly uniform in a population of cells. To test whether the level of activation is equally homogeneous, we measured the fluorescence of individual P*lrgA-gfp* cells extracted from a static, bulk culture at different times during growth. We grew cultures anaerobically in defined medium prepared with 15 mM (initial) glucose, withdrew cells periodically, dispersed them on a glass slide, and imaged them in phase contrast and GFP fluorescence on an inverted microscope. Figure 4A shows that cells showed very little fluorescence through exponential phase, up through about eight hours. At nine hours, as the cells entered stationary phase, pronounced *lrgA* reporter fluorescence was observed (Figure 4B).

**Figure 4:**
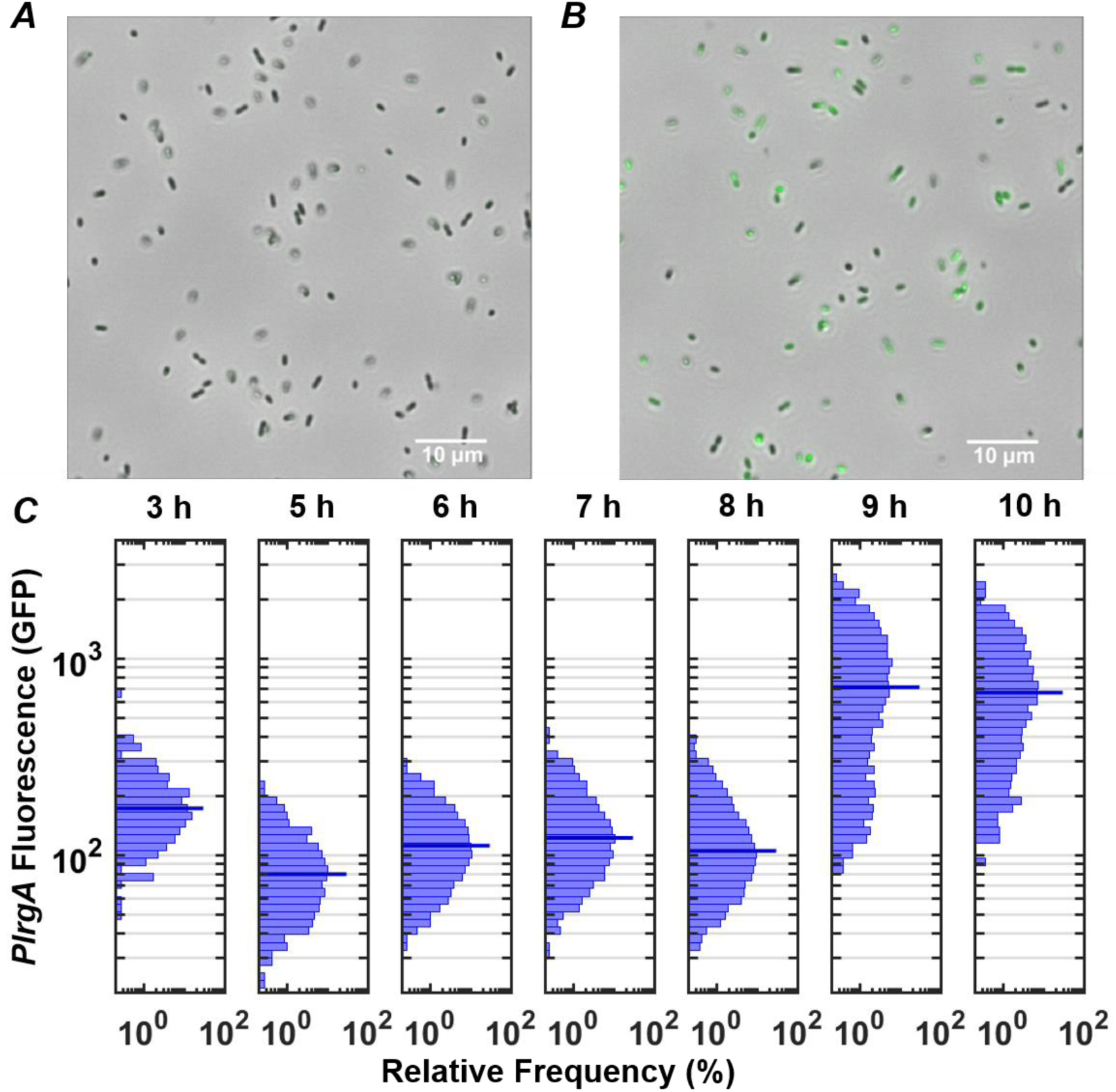
Observation of P*lrgA*-*gfp* reporter activity in individual cells extracted from bulk, anaerobic cultures grown in 15 mM initial glucose. Microscopy images of the reporter strain in phase-contrast (gray scale) are overlaid with GFP fluorescence (green) images at (**A**) 8 h and (**B**) 9 h of growth. (**C**) Histograms of individual cell GFP fluorescence measured at different times during growth. Fluorescence per cell is measured as described in (42). The length of each horizontal bar indicates the percentage of cells that fluoresced at the indicated level. The heavy horizontal line in each histogram indicates the median fluorescence of the population.

The GFP fluorescence after activation was highly variable from cell to cell, as shown by the histograms of individual cell GFP fluorescence in Figure 4C. While the histograms remain generally similar through exponential phase (roughly 3 to 8 hours following inoculation), the heterogeneity in *lrgA* activation at 9 hours is substantially greater. The median cell fluorescence at 9 h is roughly 7-fold greater than at 8 h, while the brightest cells at 9 h are roughly 10-fold brighter than the brightest cells at 8 h. The 9 h distribution has a slightly double-peaked (bimodal) character, suggesting that a subpopulation of cells have activated P*lrgA* while other cells have not. The distribution narrows only slightly by 10 h, indicating that GFP concentrations in the population change little during stationary phase. This finding is consistent with Figures 1D and 1F, where the burst of *lrgA* expression lasts less than one hour. Although the tight temporal synchrony of *lrgA* expression suggests that a single external cue triggers *lrgA* throughout the culture, the population variability in the resulting level of *lrg* expression indicates that not all cells in the static culture were immediately induced, or that the *lrgAB* operon is not so tightly regulated as to enforce a consistent response among cells once induced.

### Activation of lrgA in controlled flow requires pyruvate and deoxygenation

The observation that high initial glucose concentrations suppress the activation of *lrgAB* at the onset of stationary phase suggests that the *lrgAB* expression burst may be triggered by the exhaustion of glucose from the growth medium and the alleviation of catabolite repression of *lrgAB*. However, Figure 2 and Figure 3 also show a role for molecular oxygen, possibly in combination with extracellular pyruvate. A difficulty with using bulk, static cultures to study how these inputs affect *lrgA* is that they are altered by the growth and maturation of the culture; once a static culture is allowed to grow to stationary phase, the chemical environment of the cells is poorly defined. To identify more precisely the factors that trigger *lrgAB* we used microfluidic flow channels to apply a stable flow of fresh, defined medium to cells that were under continuous observation. We loaded P*lrgA-gfp* cells into microfluidic flow channels (*Methods*) on a microscope stage and supplied a continuous flow of fresh, defined medium through each channel. The flow rate of 20 μl/h ensured that the 1.7 μl volume of medium within each channel was replaced every 5.1 minutes. This flow prevents the cells adhered in the channels from modifying their chemical environment.

Figure 5A shows the response of cells that were provided an air-equilibrated (aerobic) defined medium containing 5 mM glucose and 10 mM pyruvate. Expression of *lrgAB* remained at basal levels throughout the experiment. (A modest decline in the average fluorescence at 180 and 210 minutes is an artifact of rampant growth affecting the image analysis algorithm). Similar flow experiments using medium that was either fully aerated, or partially deoxygenated by stirring in vacuum or under N2, produced GFP histograms very similar to Figure 5A (data not shown): No activation of *lrgA* was observed in flow experiments at any combination of glucose and/or pyruvate concentrations when the supplied media were aerobic or partially deoxygenated.

**Figure 5:**
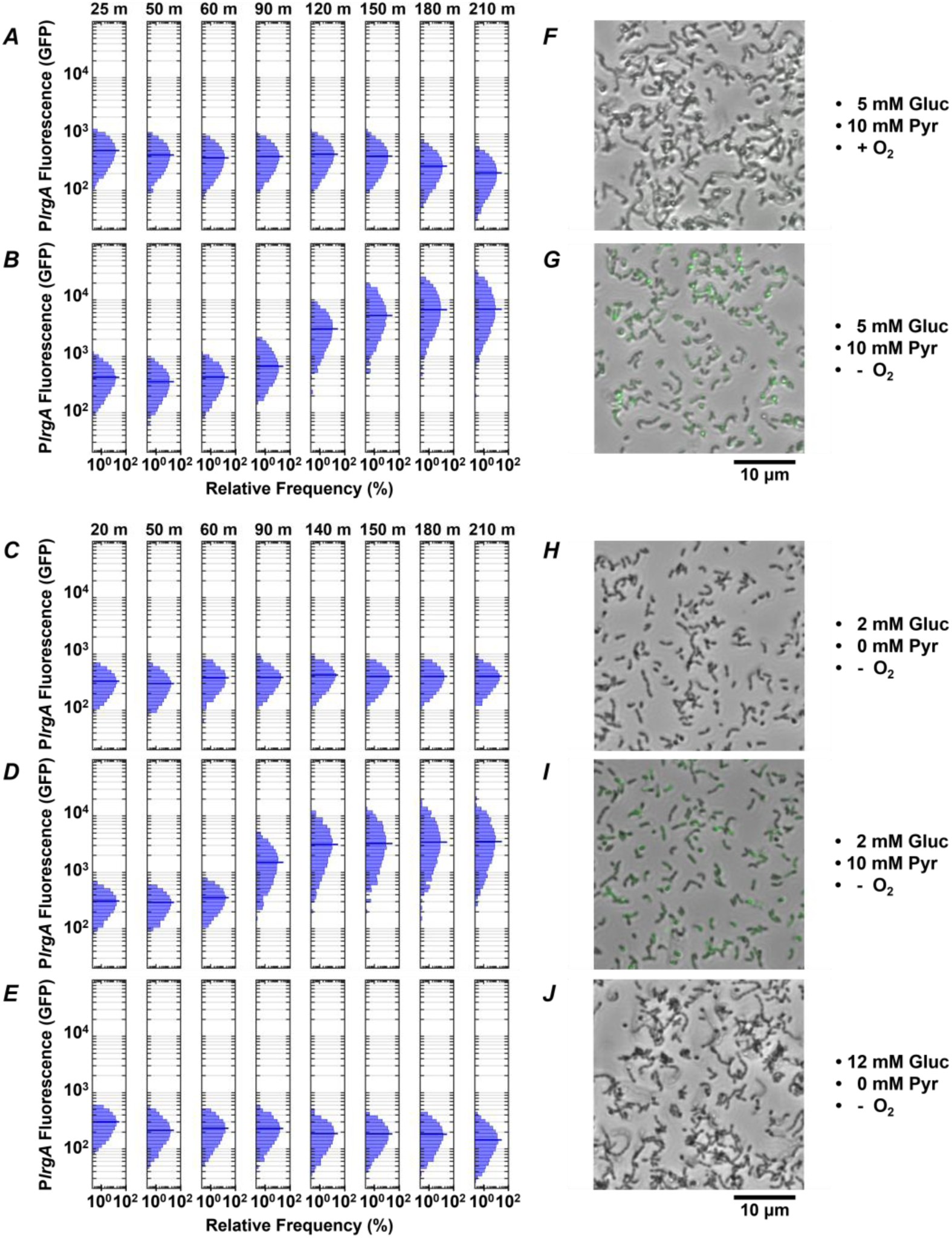
Effect of O2, glucose and pyruvate on a P*lrgA-gfp* reporter in flowing medium. The histograms show the green fluorescence of individual cells adhered within microfluidic channels and subject to a steady flow of fresh, defined medium: (**A**) aerobic medium containing 5 mM glucose / 10 mM pyruvate; (**B**) anoxic medium containing 5 mM glucose / 10 mM pyruvate; (**C**) anoxic medium containing 2 mM glucose (no added pyruvate); (**D**) anoxic medium containing 2 mM glucose / 10 mM pyruvate; (**E**) anoxic medium containing 12 mM glucose (no added pyruvate). (**F-J**) Phase microscopy images (collected at 150 minutes) of the reporter strain are shown in gray scale, overlaid with GFP fluorescence (green) images.

We therefore tested whether more rigorous deoxygenation was needed to mimic the conditions of a static, anaerobic (mineral oil layer) well plate and induce a response from *lrgAB*. Figures 5B and 5D show the results when the growth medium was made stringently anoxic by the addition of an enzymatic system that scavenges molecular oxygen (*Methods*). These highly anoxic media induced robust expression of *lrgAB*: Strong GFP production was observed after 90-120 minutes of flow of anoxic medium that contained 5 mM glucose and 10 mM pyruvate (Figure 5B), or 2 mM glucose / 10 mM pyruvate (Figure 5D). (The first 50 minutes of the 90-120 minute delay is attributable to replacement of partially deoxygenated medium that was initially present in the flow connections.)

These data demonstrate that rigorous deoxygenation is a condition for the *lrgAB* reporter to activate in a continuous flow experiment. We therefore tested whether pyruvate was also required. Anoxic medium containing 2 mM (Figure 5C) or 12 mM (Figure 5E) glucose, without added pyruvate, did not activate *lrgA*. In summary, strong upregulation of *lrgA* was only achieved under continuous flow conditions when the supplied medium was rigorously deoxygenated and contained added pyruvate. Once these conditions were present, the concentration of glucose (over the range 2-5 mM) had only modest additional effect on *lrgAB* activity. After 150 minutes in supplied medium, microscopy images in Figures 5G, 5I show cells with an activated *lrgAB* reporter (green), distinctly brighter than cells growing in aerobic or non-pyruvate media (Figures 5F, H, J).

The activation of *lrgAB* in the flow conditions of Figures 5B and 5D was slightly stronger (relative to baseline), and with a narrower population distribution, than in the static medium study of Figure 4C. In the deoxygenated flow study, the median brightness of activated cells was about 10-fold greater than the unactivated (baseline), whereas in a static, bulk culture the median activation was only 6-7 fold greater than the unactivated baseline. The 210 minute histograms in Figures 5B and 5D lack the very broad, heterogeneous *lrgAB* expression that is seen in the activated (9 h) cells in Figure 4C.

### Deletion of ccpA does not eliminate burst expression of lrgA

The above data strongly suggest that either molecular oxygen or glucose inhibits *lrgA* activation until the conclusion of exponential growth. Because a *cre*-site for the catabolite repressor protein CcpA was recently identified (22) in the *lrgA* promoter region, we investigated a possible role for CcpA in suppressing *lrgAB* activity. We compared expression of a P*lrgA-gfp* reporter in the wild type background and in a Δ*ccpA* strain, both growing anaerobically, for a range of glucose concentrations. Figures 6A and 6B show a similar abrupt onset of *lrgAB* expression at the beginning of stationary phase in the *ccpA* deletion. In Figure 6C the amplitude of the expression step is larger in the *ccpA* deletion than in UA159 background, where the relative effect is larger at low initial glucose levels. Therefore, although catabolite repression may partially inhibit the magnitude of the expression burst, it evidently does not control the timing of the burst.

**Figure 6:**
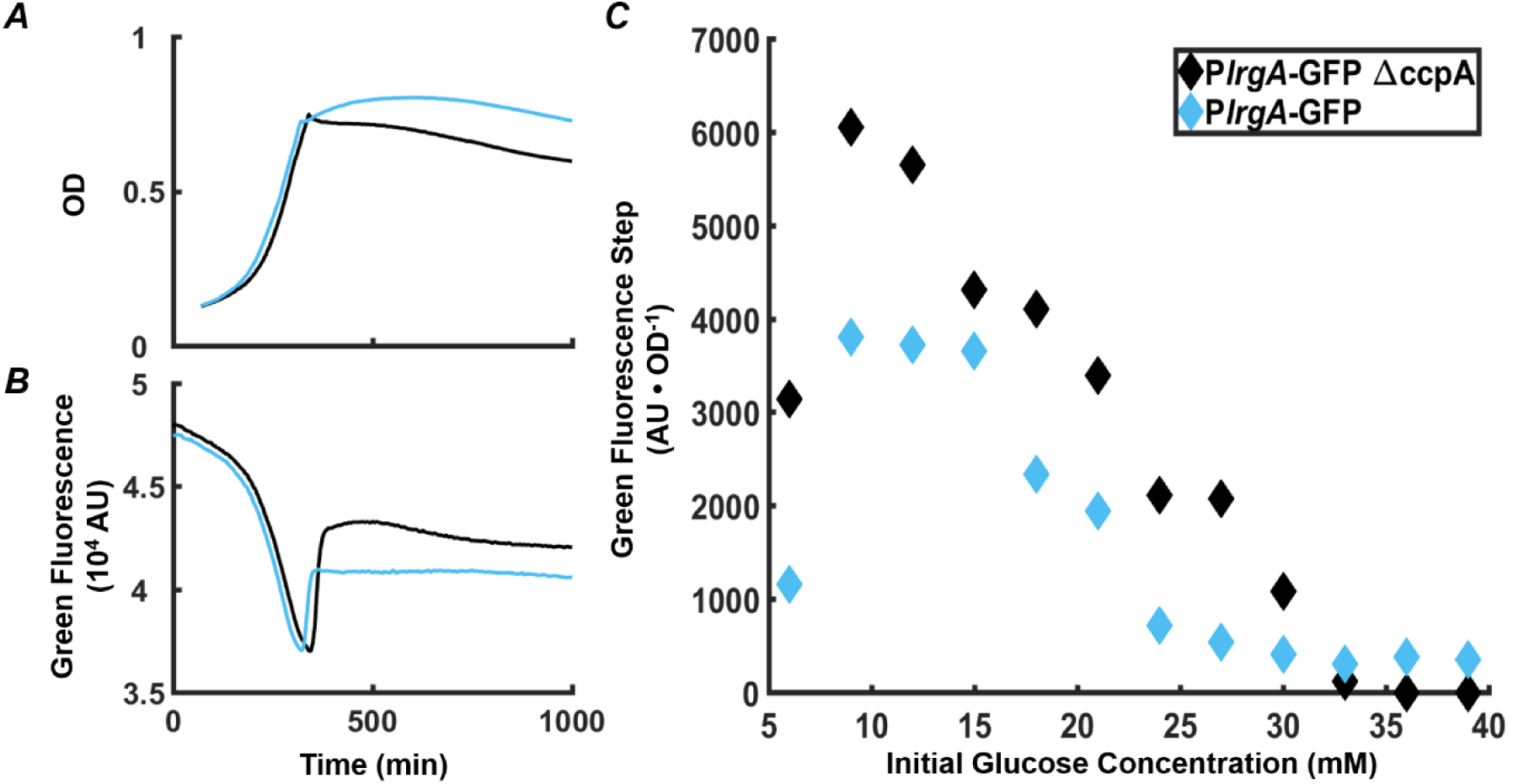
Effect of *ccpA* on the *lrgA* expression burst at stationary phase in static grown cultures. Growth (**A**) and *PlrgA-gfp* reporter fluorescence (**B**) of UA159 background (blue) and Δ*ccpA* (black) growing anaerobically in 9 mM initial glucose. (**C**) Size of the fluorescence activation step at stationary phase, normalized to optical density.

### Overexpression of lytST permits lrgA expression in aerobic media

The LytST two component system is implicated in the regulation of *lrgAB* homologs, as for example in *B. subtilis* where *lytST* was linked to pyruvate sensing and shown to be required for expression of the *lrgA* homolog (8). Prior studies of *S. mutans* in static, bulk cultures showed that deletion of *lytS* (17) or *lytST* (11) abolished the stationary phase expression of *lrgAB*. Therefore, we did not attempt to activate *lrgAB* in a *lytST* deletion strain under microfluidic conditions. However, we did test whether overexpression of *lytST* affects the expression of *lrgAB* under microfluidic flow.

We loaded a *lytST* overexpression strain harboring the P*lrgA-gfp* reporter into microfluidic channels as above. Figure 7A shows the response of cells that were provided aerobic (air-equilibrated) defined medium containing 2 mM glucose and 10 mM pyruvate. Expression of *lrgA* remained constant throughout the experiment but with a median fluorescence nearly 1.7 to 3.4 fold greater than wild type cells in a similar but deoxygenated medium (in Figures 5B and 5D). Therefore, the overexpression of *lytST* bypasses the *lrgA* requirement for rigorous deoxygenation.

**Figure 7:**
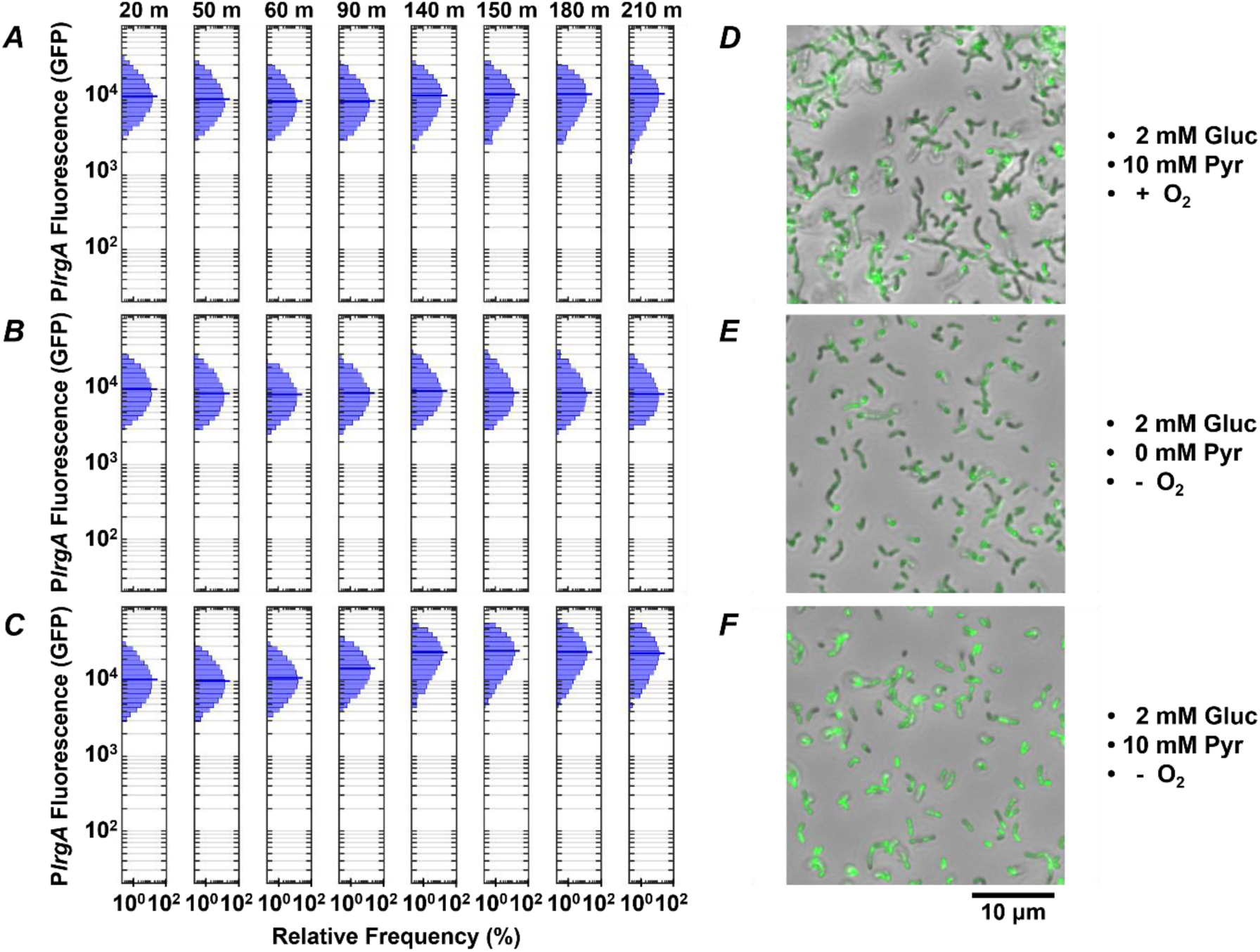
Effect of *lytST* on activation of a P*lrgA-gfp* reporter in flowing medium of fixed composition. The histograms show the green fluorescence of individual cells adhered within a microfluidic channel and subject to a steady flow of fresh, defined medium. Conditions are (**A**) aerobic medium containing 2 mM glucose / 10 mM pyruvate; (**B**) anoxic medium containing 2 mM glucose (no added pyruvate); (**C**) anoxic medium containing 2 mM glucose / 10 mM pyruvate. In (**D-F**) microscopy images of the *lytST*-overexpressing strain with P*lrgA*-*gfp* in phase-contrast (gray scale) are overlaid with GFP fluorescence (green) images collected at 150 minutes.

We tested whether pyruvate was needed to activate the *lrgA* reporter in the *lytST* overexpressing strain. Figure 7B shows that *lrgA* activated in anoxic medium containing 2 mM glucose lacking added pyruvate. Robust expression of *lrgA* was nearly identical to Figure 7A. We also tested activation of *lrgA* in anoxic medium with 2 mM glucose and 10 mM added pyruvate (Figure 7C) which were necessary to activate *lrgA* in the UA159 background. After 90-140 minutes of flow, expression of *lrgA* increased to about 2-fold greater than in Figures 7A and 7B. Figures 7D-7F show cells with activated *lrgA* reporters (green). These data show that although *lytST* overexpression does alleviate the requirement for anoxic conditions in activating *lrgA*, it does not entirely eliminate sensitivity to external pyruvate.

Finally, the population distribution of individual cell fluorescence in the *lytST* overexpression strain was observed to be slightly narrower than in the UA159 background, Figures 5B and 5D.

## Discussion

The *cidAB* and *lrgAB* operons were first identified as a putative holin-antiholin system in *Staphylococcus aureus*, with gene products that control extracellular murein hydrolase activity (3–5, 7). The *S. aureus lrgAB* operon is activated differentially through the growth curve, with the largest number of RNA transcripts detected during the transition from exponential to stationary phase (3, 7). Studies of *S. mutans lrgAB* have found generally similar patterns of expression (11, 22), although these transcriptional studies have not yielded a precise determination of the environmental cues that control the operon. By combining a fluorescent gene reporter for *lrgA* with single cell observations and microfluidic control of growth media conditions, we obtained a more detailed understanding of the environmental signals that trigger *lrgAB* in early stationary phase.

Several previous studies (11, 22) showed that higher glucose concentrations suppress *lrgAB* expression, and a recent study found a binding site for the catabolite repressor protein CcpA on the *lrgA* promoter region (22). The fact that *lrgA*, like many other virulence-linked genes in *S. mutans*, is regulated by catabolite repression via CcpA (33) could potentially explain the burst of *lrgA* expression at the end of exponential growth. However, our data imply that a different input must play the dominant role in suppressing *lrgA* during the exponential phase. Deletion of *ccpA* did not affect the timing of the expression burst (Figure 6), and it had only a modest, qualitative effect on the level of that expression, increasing it less than two fold (18).

By contrast, molecular oxygen was found to exert decisive control over the *lrgA* burst. No combination of glucose/pyruvate concentrations was found to activate *lrgA* in cells that were growing in a continuous flow of fresh, defined medium, unless that medium was rigorously deoxygenated. In deoxygenated medium, robust *lrgA* expression occurred even though the composition of the medium (FMC medium containing added pyruvate) was otherwise compatible with normal, exponential growth. This finding suggests that the population-wide, tightly synchronized burst of *lrgA* expression observed in static, bulk cultures at stationary phase is not triggered by an internal state of the bacteria, or by accumulation of pyruvate or depletion of nutrients from the media, but rather by exhaustion of molecular oxygen. Exhaustion of oxygen is presumably an all-or-nothing signal that occurs at a well-defined time point during the growth curve, unlike the gradual accumulation of a waste product or a quorum sensing signal.

A previous *S. mutans* study reported that *lrgAB* was upregulated when grown aerobically in static, bulk cultures (2). That study used a microarray analysis to compare total RNA between aerobic and anaerobic cultures of *S. mutans* during mid-exponential phase (optical density of 0.4 at 600 nm) (2). A followup study similarly reported that *lrgAB* expression at stationary phase was much more pronounced when grown aerobically compared to low-oxygen growth (17). Our present study differs from these two earlier studies in some key respects. One is that our use of a fluorescent reporter allows us to characterize the large burst of *lrgAB* activity that occurs within a very narrow temporal window at stationary phase, which may be missed in the transcriptional study. Oxygen concentration could potentially also affect RNA stability. In addition, the low-oxygen conditions in (2) and (17) are less well defined than in the present study. For example the low-oxygen condition in (17) consisted of growth in 5% CO2, which is not equivalent to the stringently anaerobic condition achieved here through the use of an enzymatic oxygen scavenger. Our data clearly show that a high level of control over oxygen concentration, in addition to high time resolution, are both necessary in order to characterize *lrgA* activity early in stationary phase.

The mechanism by which oxygen represses *lrgAB* is not known, although the VicRK two component system is a potential candidate that has been shown to influence *lrgAB* expression (18). VicRK has also been linked to oxidative stress tolerance in *S. mutans* (24, 25, 34) and VicK is regarded as a potential sensor of oxygen or redox conditions (35). Consistent with this interpretation, a transcriptional study found that deletion of *vicK* led to moderate increase in exponential phase expression of *lrgA*, but a nearly 100-fold decrease in stationary phase expression (18).

LytST has also been identified as a potential intermediate between molecular oxygen and *lrgA* (11, 17). However, LytST homologs in other organisms have recently been identified as sensors of extracellular pyruvate. In *E. coli* two component systems of the LytS/LytTR family have been identified as a receptors for external pyruvate (36, 37), and a LytST-regulated system is triggered by extracellular pyruvate (32). In *B. subtilis* both *lytST* and the *lrgA* homologs, *ysbA* and *pftA*, were shown to be essential for pyruvate utilization (8). As in *S. mutans*, *B. subtilis ysbA* activates at the onset of stationary phase and decreases its expression with increasing initial glucose concentrations due to regulation by CcpA (8, 9). The *ysbAB* (or *pftAB*) operon is induced by LytST in the presence of extracellular pyruvate (9). That study reported that PftA and PftB form a hetero-oligomer that functions as a pyruvate-specific facilitated transporter and, together with LytST, help to adapt to a changing environment when the preferred carbon sources have been exhausted (9).

Certainly the LytST system is a key regulatory input to *lrgAB* expression in *S. mutans*, as deletion of *lytST* was previously shown to prevent stationary phase expression of *lrgAB* (11). In our studies a *lytST* overexpressing strain readily activated *lrgA*, even in the absence of pyruvate and in media that were not thoroughly deoxygenated. LytST overexpression eliminated the bursting character of *lrgA* expression and caused instead generally robust expression under aerobic and pyruvate-absent conditions where *lrgA* expression was absent in the wild type. These data indicate that *lytST* is not only required for activation of *lrgA*, but that it can overpower the repression signals due to molecular oxygen. Interaction of LytST with the *cre1* site previously suggested that LytST may inhibit the action of CcpA, and therefore partially bypass catabolite repression as well (22).

The very brief duration of the *lrgA* expression burst at stationary phase may offer an intriguing clue to the mechanisms of its regulation, as it suggests a self-limiting behavior. In the *B. subtilis* study above, induction of the *lrgAB* homolog *ysbAB* (*pftAB*) increased as pyruvate increased up to 1 mM but was also inhibited via LytST under excess pyruvate conditions (9), suggesting that an influx of pyruvate led to inhibition. One may speculate that if expression of *S. mutans lrgA* triggers a pyruvate influx that suppresses further *lrgA* expression, then the temporal profile of *lrgA* activity in response to extracellular pyruvate would appear as a rapid burst as is observed here. In that case, if pyruvate can also enter the cell by another pathway (unrelated to LrgAB and LytST), then very high concentrations of extracellular pyruvate would be expected to suppress *lrgAB* activity, as is observed. A pyruvate-dependent self-limiting mechanism of this type is consistent with findings that overexpression of *lrgAB* from a plasmid led to upregulation of *lrgAB* during exponential phase but downregulation of *lrgAB* during stationary phase (18).

It is an interesting property of *lrgAB* that its activation (and subsequent deactivation) in a bulk culture is tightly synchronized temporally in the population, and yet the level of expression (as indicated by GFP concentration) is variable in individual cells. Although some of the cell-to-cell heterogeneity seen in *S. mutans* fluorescent protein expression can probably be attributed to the use of plasmid-based reporters (38) the heterogeneity we observe in *lrgA* expression cannot be due entirely to the plasmid reporter. When cells drawn from a static culture activate *lrgA*, the population distribution in fluorescence is broad with a strongly bimodal character (Figure 4C). This bimodality is highlighted in Figures 8A and 8B, which represent each of the *lrgA-*active, single-cell histograms as the sum of two gamma probability distributions. (The gamma distribution is characteristic of stochastic gene expression (39)) The relative areas under the two distributions indicate that roughly 82% of cells are *lrgA-*active (high fluorescence) at 9 h, and about 95% are *lrgA-*active at 10 h. By contrast, *lrgA* expression under microfluidic flow (Figures 5B, 5D) lacks this bimodal character, producing virtually unimodal histograms (≥ 98% *lrgA-*active) in the same mathematical representation (Figure 8C, 8D). This finding indicates that, when environment conditions are sufficiently uniform as in the microfluidic study, a robust and generally similar level of *lrgA* expression is observed population-wide. Therefore, local differences or gradients in key parameters such as pyruvate, oxygen or glucose may explain some of the heterogeneity that was observed in our static culture studies, and also in the individual cell expression of the *lrgA* homolog *ysbA* in *B. subtilis* (8).

**Figure 8:**
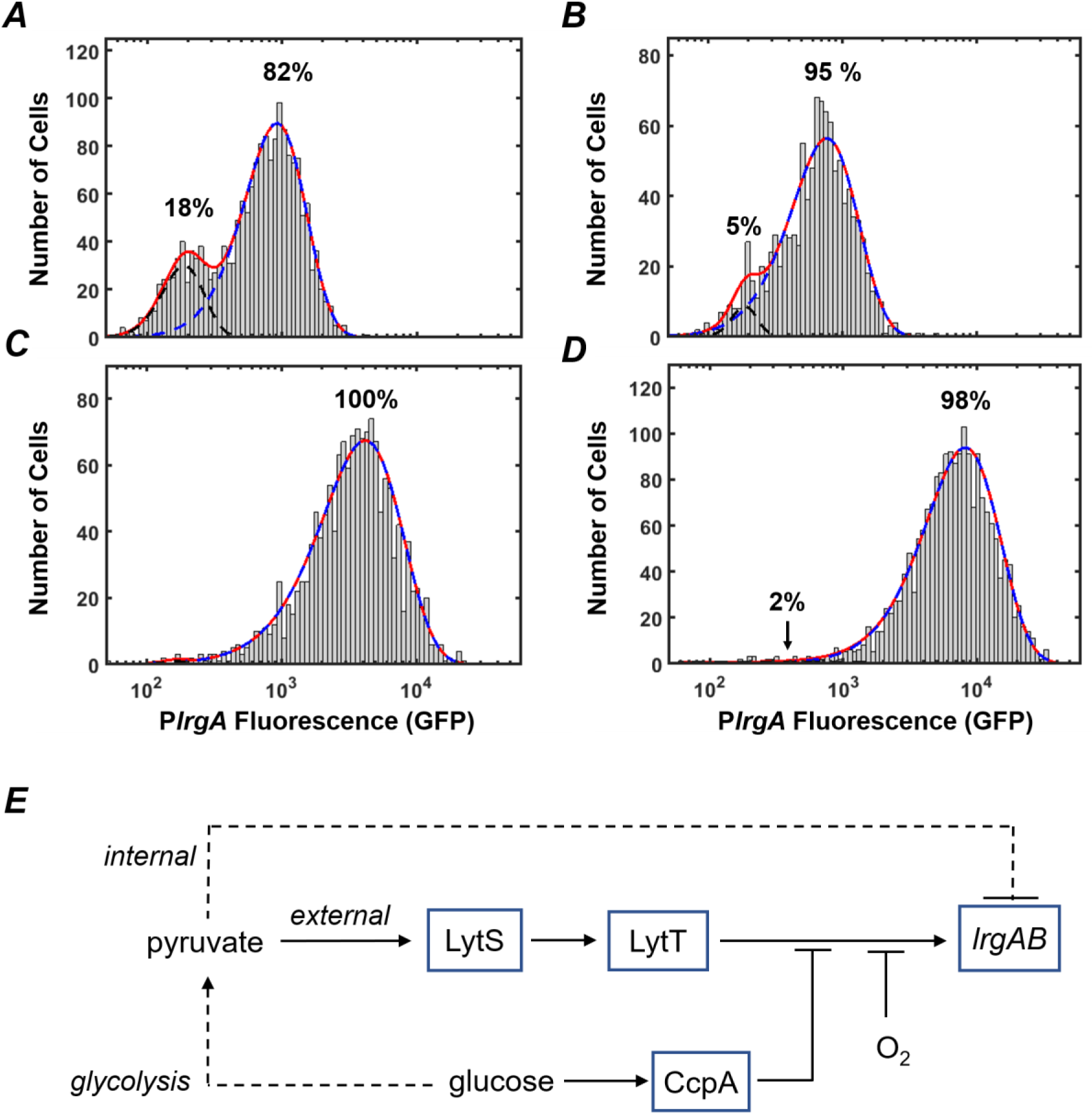
P*lrgA-gfp* reporter activity of individual cells fit to two gamma distributions representing the activated (dashed blue) and inactivated (dashed black) populations. Solid red line is the sum of both distributions. (**A**) 9 hour sample from Figure 4C; (**B**) 10 hour sample from Figure 4C; (**C**) 210 minute sample from Figure 5B; (**D**) 210 minute sample from Figure 5D. (**E**) Schematic for control of *S. mutans lrgAB* by glucose, molecular oxygen, and external and internal pyruvate signals, via LytST.

The presence of heterogeneity without bimodality in our microfluidic data also implies that *lrgAB* is regulated in an open-loop mechanism, without benefit of the positive transcriptional feedback that is typically associated with bimodality in gene expression (40). Rather, a mechanism of activation by LytST followed by negative feedback via intracellular pyruvate, as hypothesized above (Figure 8E), may be sufficient to control *lrgAB*, as it allows both an on-switch and an off-switch. We note that the histogram of single-cell fluorescence is markedly narrower for the *lytST* overexpressing strain (Figure 7A) than for the wild type background (Figure 5D), suggesting that *lytST* overexpression is a strong enough stimulus that it brings *lrgA* expression closer to saturation and reduces the heterogeneity that is normally present.

Finally, our study has not identified a pathway by which *cidAB* modulates *lrgAB* expression. These two operons exhibit a complex pattern of transcriptional cross regulation that is growth phase dependent, indicative of interactions between different gene products within both operons. It is likely not as simple as mutual repression (18). Future studies of *cidAB* activation may begin to shed light on how the two operons interact.

## Acknowledgments

The authors acknowledge funding support from the NIDCR through awards R01 DE025237 and R01 DE023339.

## Methods

### Bacterial strains, plasmids and growth conditions

Observing the effects of pyruvate and glucose on *lrgA* in *S. mutans* was possible through a *gfp* fusion to the promoter region of *lrgAB*, which was inserted into the pDL278 shuttle vector (carrying spectinomycin resistance) as described in (22). The resulting plasmid was inserted into a wild-type UA159 and a *ccpA*-deficient mutant (41) to give the P*lrgA-gfp* and Δ*ccpA*/P*lrgA-gfp* strains, respectively (22).

The *lytST* overexpression strain was constructed using the method described in (18). Briefly, a fragment containing the *ldh* promoter region (P*ldh*) and a polar kanamycin resistance gene (ΩKm-P*ldh*) was used to replace the *lytST* promoter region: two 0.5 kb fragments surrounding the −35 and −10 regions of the *lytST* promoter were amplified and ligated to the ΩKm-*Pldh* cassette and then transformed into *S. mutans*.

For studies of P*lrgA* activation in a well plate system, overnight cultures of *S. mutans* UA159 and its derivatives were incubated in complex medium BHI with 1 mg/ml spectinomycin to ensure plasmid homology at a temperature of 37 °C in an atmosphere composed of 5% CO2. Overnight cultures were washed twice in phosphate buffered saline (PBS) of pH 7.2. They were then diluted 1:100 into defined medium (FMC) pH corrected to 7.0 containing final concentrations of glucose and pyruvate dictated by the experiment conducted. Fresh cultures were allowed to grow to early exponential phase with an OD600 of 0.1 before being followed by any further testing. For single cell studies as well as studies under a flow environment, overnight cultures of *S. mutans* were grown in BHI supplemented with an additional 20 mM glucose to ensure no activation of *lrgA*. Overnight cultures were washed twice in PBS and diluted 1:35 in fresh FMC before allowed to incubate to an OD600 of 0.1.

### Measuring growth and gene activation in bulk

The data seen in Figures 1 – 3, 6 was measured using a BioTek Synergy 2 multimode plate reader. Overnight samples were first diluted 100-fold into fresh FMC media with the prepared initial carbohydrates necessary for the experiment. Samples were grown to an OD600 of 0.1 in prepared FMC media before being dispersed into 2 ml volumes (Figures 1,2,6) or 200 μl (Figure 3) on 24 or 96 well plates respectively. Samples were covered with a 410 μl mineral oil overlay to facilitate anaerobic growth on a 24 well plate and 75 μl on a 96 well plate. Aerobic growth was facilitated with no mineral oil overlay and the plate was set to shake for 10 seconds every two minutes. Cultures grew in the well plates for 24-35 h to reach well into stationary phase and its growth was monitored by its optical density at 620 nm which was measured at 5 minute intervals. Fluorescence was monitored by a green filter at 485-520 nm.

### Measuring lrgA activation from bulk

The fluorescence increase seen at the onset of stationary phase was calculated by calculating the time derivative (slope) of the fluorescence curve obtained from the BioTek Synergy 2 and finding the time value at maximum slope. This time value corresponds to the inflection point of the fluorescence increase. An adjacent local minimum and maximum in the fluorescence are then found from the nearby time values at which the time derivative crosses zero. The difference between these maximum and minimum values is the fluorescence step at the onset of stationary phase. We then normalized this fluorescence step, dividing it by the optical density of the culture at its entry into stationary phase.

### Slide Experiments

Overnight cultures were diluted 1:35 fold into a 20 ml seed culture with an oil overlay inside an incubator maintaining a 5% CO2 atmosphere at 37 °C. To take phase and fluorescence images, a 600 μl sample was collected into a cuvette from the seed culture and an OD600 measurement was taken. The same sample was then ultra-sonicated to break up the cell chains and 4 μl deposited on a glass coverslip. Phase contrast and fluorescence images of the slide were taken on a Nikon, TE2000U, inverted microscope together with a Photometric Prime camera and a green filter. Phase and fluorescence images were taken periodically throughout the full growth cycle of the culture until a stable, stationary phase optical density was reached. GFP concentration of individual cells was assessed from microscopy images using a method described previously (42).

### Microfluidic Design

An ibidi μ-slide VI (ibidi USA, Inc) was used to measure activation levels of PlrgA under flow of medium at set rates. The ibidi slide consisted of 6 flow channels that had dimensions of 0.1 × 1 × 17 mm for a total volume of 1.7 μl. Each of these rectangular channels had allowed viewing through a microscope. Each channel had an inlet and an outlet that fit a standard luer fitting which allowed solutions of the desired media to be pumped through the flow channels. The ibidi μ-slide was secured to the stage of a Nikon, TE2000U, inverted microscope that is housed inside a temperature controlled Lexon chamber. While data was collected, the chamber was maintained at a constant 37 °C by an electronic temperature controller.

### Microfluidic Experiments

We cultured *S. mutans* P*lrgA-gfp* cells in defined (FMC) medium containing initially 10 mM glucose and grew them to 0.3-0.4 OD. We then sonicated the cells to break apart chains and loaded the cells into microfluidic flow channels (1.7 uL volume per flow channel, 100 μm channel depth, six independent channels per flow device, Ibidi GmbH). Cells were allowed to settle onto the lower window of the channel for 20 minutes, while the channel was mounted onto an inverted microscope in a temperature-controlled chamber. A flow of fresh medium was then supplied into the channels by a syringe pump at a rate of 1000 μl/h for 30 minutes to replace and refresh the medium in the channels, connections and fittings. After the 30 minute purge, the pump rate was reduced to 20 μl/h and held constant for the duration of the experiment.

To ensure that the growth media for the microfluidic studies was fully deoxygenated, we added an oxygen scavenging system consisting of 2 U/ml glucose oxidase and 120 U/ml catalase (43). This enzymatic system rapidly consumes O2 from the medium by breaking down glucose to yield gluconic acid and H2O as products. Although the glucose oxidase generates H2O2 as an intermediate product (which is then broken down by the catalase), *S. mutans* is tolerant of low to moderate concentrations of H2O2 far higher than would be present during this reaction (31).

